# Embeddings of genomic region sets capture rich biological associations in lower dimensions

**DOI:** 10.1101/2021.05.07.443166

**Authors:** Erfaneh Gharavi, Aaron Gu, Guangtao Zheng, Jason P. Smith, Aidong Zhang, Donald E. Brown, Nathan C. Sheffield

## Abstract

**Motivation:** Genomic region sets summarize functional genomics data and define locations of interest in the genome such as regulatory regions or transcription factor binding sites. The number of publicly available region sets has increased dramatically, leading to challenges in data analysis.

**Results:** We propose a new method to represent genomic region sets as vectors, or embeddings, using an adapted word2vec approach. We compared our approach to two simpler methods based on interval unions or term frequency-inverse document frequency and evaluated the methods in three ways: First, by classifying the cell line, antibody, or tissue type of the region set; second, by assessing whether similarity among embeddings can reflect simulated random perturbations of genomic regions; and third, by testing robustness of the proposed representations to different signal thresholds for calling peaks. Our word2vec-based region set embeddings reduce dimensionality from more than a hundred thousand to 100 without significant loss in classification performance. The vector representation could identify cell line, antibody, and tissue type with over 90% accuracy. We also found that the vectors could quantitatively summarize simulated random perturbations to region sets and are more robust to subsampling the data derived from different peak calling thresholds. Our evaluations demonstrate that the vectors retain useful biological information in relatively lower-dimensional spaces. We propose that vector representation of region sets is a promising approach for efficient analysis of genomic region data.

**Availability:** https://github.com/databio/regionset-embedding

## Introduction

An epigenomics experiment is often represented as a region set, which is a collection of genomic intervals that identify the locations of interest produced by a biological experiment. Region sets are produced from various experiments, such as ChIP-Seq^1^ or ATAC-Seq^2,3^, and contain the locations of functional elements along the genome such as enhancers, promoters, and transcription factor binding sites^4^. Region sets are frequently stored as Browser Extensible Data (BED) files, which may contain up to millions of individual regions. As the amount of publicly available epigenome data has increased, the volume of data has led to challenges analyzing it.

In the past few years, many methods have been developed for processing, analyzing, and comparing genomic region sets. One key task has been finding connections between region sets, but like many other tasks, this is complicated by the volume and complexity of region set data^5–9^. Here, we address this problem by embedding regions sets in a lower dimensional space that retains important biological information. Robust embeddings have potential to highlight important biological relationships among region sets and can lead to new ways to query and analyze data repositories. Importantly, robust embeddings can capture biological information that would be difficult to extract from raw data, and therefore the performance of downstream tasks, such as classification and clustering, can be improved^10^.

One simple example of data representation is the bag-of-words method for textual data, which uses the existence of words to create a binary vector representation. More recently, new methods have invoked the distributional hypothesis, which states that words in similar context have similar meanings^11–13^. In this framework, embeddings learned from context have led to improved performance in a wide range of natural language processing (NLP) tasks^14^, and distributed representations of data are now commonly used across many disciplines to reduce data dimensionality and learn relationships. In bioinformatics, embeddings have been used to represent DNA nucleotide sequences^15–18^, protein sequences^19^, genes^20^, and single-cell Hi-C data^21^, among others. Factorized tensor decomposition has also been used to achieve biologically meaningful representation for RNA-Seq data^22^. Most similar to our use case is the recently published Avocado method, which learns a dense representation for a region using a deep neural network trained on epigenome signal information^23^. Avocado uses metadata annotations during the training to develop the embeddings, which are then evaluated on other downstream tasks. Our approach differs in several ways: first, it is more general in that it trains on intervals only and does not require signal data, which makes it amenable to a larger class of data that may or may not have accompanying signal annotation (e.g. CpG island annotation, HMM chromatin states, k-mer locations, motif matches, etc); second, training is based solely on co-occurrence of intervals without requiring metadata annotations; third, the underlying model uses the shallow word2vec neural network; and fourth, our primary goal is to create embeddings for *sets* of regions, whereas Avocado is geared toward representations of individual regions. Therefore, while there is clear value in methods that incorporate signal value and annotation information, there is also utility in the more general approach we present here.

We translate NLP techniques to genomic region set data by considering each region set as a text document, with each region as a word inside that document. We applied two NLP approaches in our work: a method based on term frequency-inverse document frequency^24^ for feature selection, and word2vec embeddings (introduced in Supplemental Materials)^11^. The proposed condensed representation for a region set is learned using only genomic coordinates. For comparison, we also tested a non-condensed representation based on a simple binary vector approach. We evaluate with three evaluation tasks: first, a classification task to predict and visualize biological characteristics of a region set, such as cell line, antibody, and tissue type; second, a similarity detection task to assess if the models capture differences between a reference file and simulated perturbed ones; and third, a subsampling task to test robustness to data mimicking varying thresholds for calling peaks. Using these evaluations, we show that the proposed embeddings maintain high classification performance, quantitative reflection of simulated perturbation, and robustness to subsampling by different peak calling thresholds, despite multi-fold reduction in dimensionality.

## Materials and methods

### Overview of the approach

We used a dataset from the ChIP-Atlas database, which contains uniformly processed ChIP-seq data from the Sequence Read Archive (Fig. 1a)^25^. We applied three different approaches to represent these region sets: union representation, tf_idf-based representation, and region-set2vec embedding (Fig. 1b). To evaluate the embeddings, we employed three machine learning tasks: classification, similarity detection, and peak threshold robustness (Fig. 1c).

**Figure 1:**
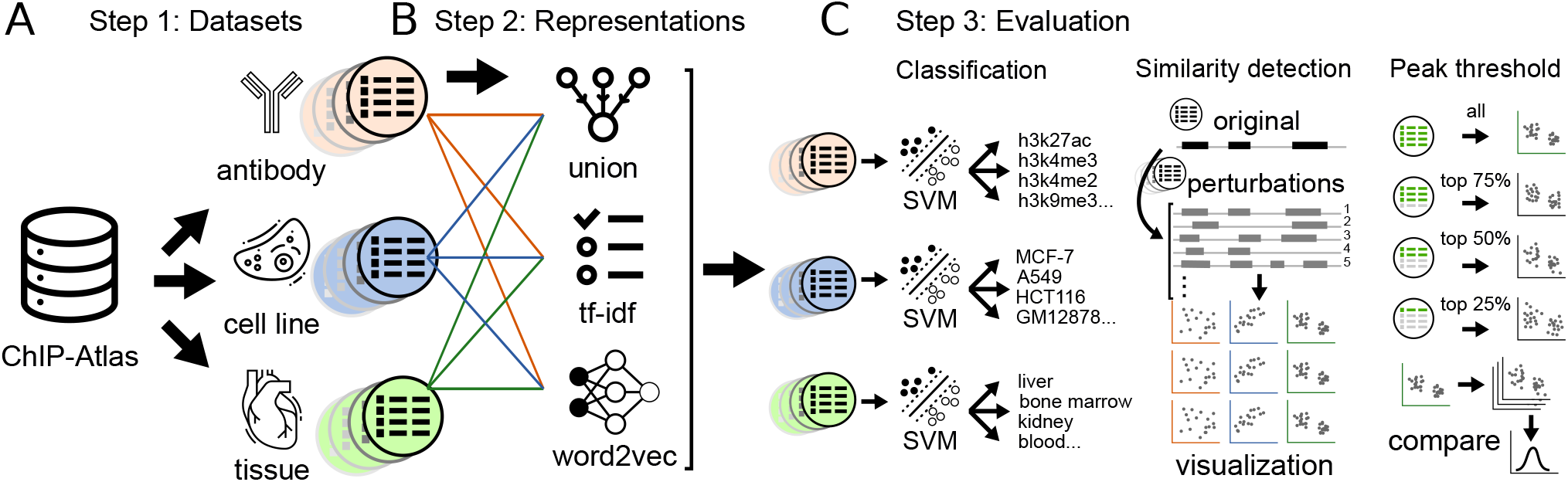
Overview of methods. (**a**). Datasets were divided into three different experiment types from the ChIP-Atlas dataset. (**b**). Each file was converted into three different vector representations. (**c**). We evaluated the representations with three evaluation tasks. First, classification of the region set representations into tissue type, cell line, and antibody. Second, evaluating sensitivity to known changes in the data using a simulated dataset. Third, testing the robustness of the embeddings to different subsets of the original region list.

### Dataset

Chromatin immunoprecipitation followed by sequencing (ChIP-seq) identifies the binding sites of DNA-associated proteins. We downloaded 12,731 BED files representing ChIP-Seq data from ChIP-Atlas and constructed 3 different test datasets: one annotated with antibody, one with cell line, and one with tissue type.

The antibodies are *h3k27ac, h3k27me3, h3k4me1, h3k4me2, h3k4me3, h3k36me3*, and *h3k9me3*. The cell lines are *MCF-7, HeLa, HEK293, A549, Hep G2, HCT116, LoVo, GM12878, LNCap*, and *K562*. The tissue types are *liver, peripheral blood, primary prostate cancer, blood, breast, bone marrow*, and *kidney*. For each of these three datasets, we divided the data into a training (80%) and test (20%) set (Table 1).

**Table 1:** Dataset statistics

|  | Classes | Training samples | Test samples |
| --- | --- | --- | --- |
| antibody | 7 | 2777 | 695 |
| cell line | 10 | 5527 | 1368 |
| tissue | 7 | 1788 | 440 |

### Representation methods

For each of our 3 test datasets, we represented the data in 3 different ways:n

#### Union representation

The union representation represents a region set as a binary vector, with each position in the vector indicating whether a particular region is present in that set (Fig. 2a). The first step is to create a consensus set of regions across all region sets in a dataset, which we refer to as a *universe* of possible regions. We created a universe by first concatenating regions in 100 random training files from each dataset into one file, then merging any regions that were closer than 1000 base pairs into a larger region using the start position of the first region and the end position of the second region (Table S1). To confirm that the random selection of 100 files did not have an impact on the universe, we created multiple union representations with different universes, which resulted in similar performance (Table S2). The resulting *n* regions in the universe correspond to the vector of length *n*, with each region specifying a position, or feature, in the vector. The binary vector representation reflects, for a particular set, which of these universe regions is present in the set, which we evaluate with a simple interval overlap calculation.

**Figure 2:**
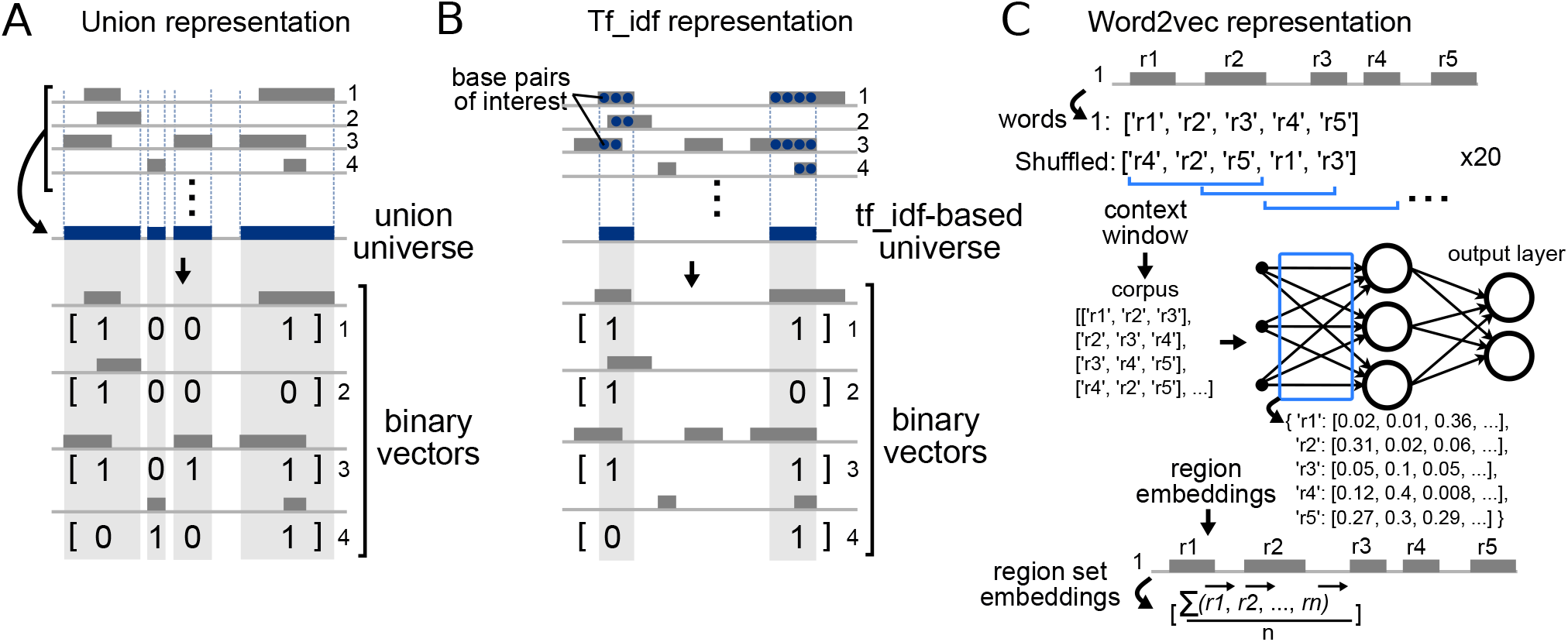
Details of embedding methods. (**a**). The union representation simply merges the region sets to create a universe, then casts each region set as a binary vector reflecting presence or absence of the merged universe region. (**b**). The tf-idf approach builds a restricted universe based on nucleotides determined to be highly informative. (**c**). The word2vec approach extracts embeddings as weights from a shallow neural network trained using co-occurrence frequencies.

#### Tf_idf-based representation

One problem with the union representation is that merging close regions creates larger regions that can obscure real biological differences in regions. To mitigate this, our second approach employs a feature selection and representation method from NLP to retain the regions that play an important role in distinction of the data (see Supplemental Material). Briefly, we consider each base pair location as a term and a region set as a document, and calculated the tf_idf score for each base pair across regions sets (Fig. 2b). Base pairs with higher scores are more informative for identifying relationships among region sets, as this approach will weight elements present in an intermediate number of region sets, while downweighting elements that are either very common or very rare across region sets.

We selected the top 100k, 500k, and 1,000k scoring base pairs (kb) for each chromosome to use in our experiment, producing three tf_idf-based representations. This was also to show the effect of the number of selected base pairs on classification accuracy. After selecting the important base pairs, we merge any adjacent base pairs to build a region and remove regions with fewer than 100 base pairs. Like the union representation approach, these regions can then be features of a binary vector. The key difference between this approach and the union representation approach is that here we are creating vector features from merged base pairs that are considered as informative.

#### Region-set2vec representation

So far, these two approaches have represented the data as high dimensional vectors. These approaches consider each region as a separate feature and do not take into account relationships among the regions. Furthermore, high dimensionality increases time and space complexity for processing and analyzing data. To address these challenges, we adapted the word2vec algorithm to train a distributed representation for each region in our training region sets (Fig. 2c). Similar to the pre-processing step of vocabulary building in NLP, we first map the regions from each file into a standard universe. We chose the union representation to be the universe because it did not use any feature selection, and is thus a fair comparison between the region-set2vec embedding and the tf_idf representation. During the training phase of word2vec, the task is to predict co-occurring regions given an input region in the same region set. After training, the trained weights in the neural network consist of 100-dimensional vector representations for each region, which we take as region embeddings.

Word2vec considers consecutive *words* in a context window (set to 100 words in our experiment) to train the embeddings. We want to capture the co-occurrence information for all the regions in a bed file. Ideally, the context window should have a length that could cover all the regions in a BED file. However, since a BED file could have thousands of regions or more, a very large context window would require very large memory and make it very hard to optimize the region embeddings. To circumvent these, we use a small constant window size at the cost of shuffling the regions multiple times. Since the order of the regions is not important, we can shuffle the regions in each region set before sampling the context window and passing them as the input of word2vec algorithm. With shuffling, all the regions that co-occur in a region set can appear in the same context window by random chance. With a smaller window size, we need less memory but a larger number of shuffling operations. Too small context window will limit the learning capability of region-set2vec, as very few regions are used at a time. In practice, we found that setting the context window size to 100 and shuffling each region set 20 times is a good choice for the current study.

After obtaining the vectors for each region, we used them to calculate a vector representing the region set by averaging all the vectors of the regions contained in the region set 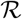, shown in Equation 1.

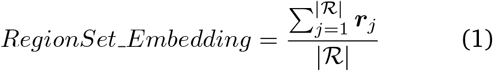

Here, 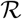 is the region set and *r_j_* is the region embedding of the *j*th region in 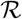.

To get a document embedding from word embeddings, a commonly used method is doc2vec^26^, which extends word2vec by introducing an additional document embedding vector in the word2vec architecture such that it can be trained jointly with word embeddings. In the context of region embedding, since the order of regions in each region set is not important and we shuffle during training, the learned document embedding can not catch meaningful information throughout the training. In fact, when we trained the doc2vec model with the same amount of data that was used for word2vec training, the performance was much lower than averaging the region embeddings (Table S3). Therefore, we find that averaging over all the region embeddings obtained from a bed file is a simple and effective document embedding method.

### Evaluation

Having created different vectors to represent region sets, we next sought to evaluate how well each proposed representation retains biological information. We designed three evaluation tasks: a classification task, a similarity detection task, and a peak threshold robustness task. The classification task asks how well the region set embeddings can reflect known biological relationships among region sets. The similarity detection task uses simulated random perturbations to assess how well the embeddings reflect known levels of mathematical difference among region sets. Finally, the peak threshold robustness task tests how much the embeddings change when the input data is truncated a subset of the regions, such as would happen if a different peak calling threshold were used.

#### Classification task

The classification task is as follows: given the region set vector representations as input, we trained classifiers to classify region sets either by tissue type, antibody, or cell line. Recall that this annotation information is not used to construct the original embeddings, which is unsupervised and relies on the regions themselves; we incorporate the annotation information in this classification task to evaluate whether the embeddings can capture the annotation information *de novo*. We employed a support vector machine (SVM)^27^ classifier for each of these tasks. An SVM is a supervised machine learning algorithm that creates a hyperplane that separates the data into sets. To classify multi-class data, we used one-vs-rest to split the data to a binary dataset for each class.

##### Evaluating classifier performance

To evaluate the performance of the classification algorithms, we used the micro-averaging of the F1 score^28^ to account for the class imbalance. This F1 score is composed of micro-precision and micro-recall, which are defined as follows:

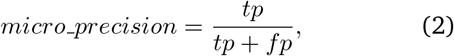

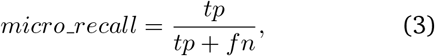

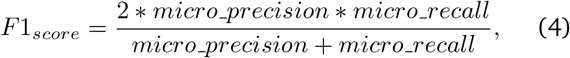

where the true positive 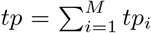 is the summation of all true positive numbers in each class, and *f_p_*, and *f_n_* are similarly defined as the summation of all false positive and all false negative numbers, respectively.

##### Visualising embeddings with UMAP

To further evaluate the representation approaches, we visualized the embeddings in 2 dimensions using uniform manifold approximation and projection (UMAP)^29^. UMAP is a nonlinear dimensionality reduction method designed to analyze high dimensional data. It has been proven as an effective tool to reveal meaningful structure in biological data^30,31^.

#### Similarity detection task

Querying and retrieving similar region sets from huge datasets relies on detecting the similarity among region sets. As mentioned, representing data in lower dimensions facilitates this process. We employ the similarity detection task to further evaluate the enrichment of our representation approaches.

##### Creating a simulated dataset

We created a simulated dataset using bedshift, a tool that randomly perturbs BED files by adding, dropping, or shifting regions to a parameterizable degree^32^. These perturbations create new files with a defined similarity that can be used as a benchmark for similarity scoring. For example, the similarity between the perturbed file and the original file should be greater if 10 percent of regions are dropped than if 20 percent of regions are dropped. The simulated dataset used all of the three perturbations, add, drop, and shift, at percentage rates from 10 to 90 percent in increments of 10 percent. For each level of perturbation, we created 100 replicates, resulting in 900 total files. We then converted the files in all datasets to the numerical representation using three methods and visualized each representation on the perturbed datasets.

##### Visualizing perturbed embeddings with UMAP

We used UMAP to plot the simulated datasets to depict the effectiveness of each representation at detecting similarities. We also did a sensitivity analysis to test the effect of three major UMAP hyperparameters: the distance metric; the number of neighbors to consider; and the minimum distance allowed for points to be in the low dimensional representation.

#### Peak thresholds subsampling task

For our final evaluation task, we asked how much the embeddings would change if trained on only a subset of the data. Genomic regions are often generated by calling peaks from the ChIP-seq signals based on a threshold. We investigate the robustness of different representations to the threshold of peak calling from signals using the classification task. Using the cell line dataset, we binned the regions based on signal values into 4 quartiles. Four datasets from the cell line annotated dataset were generated by this approach. The first is the original data that contains all of the regions. We refer to this dataset as *All*. Three other datasets were generated by selecting the top 75%, top 50%, and top 25% of the regions in each bed file based on the signal values. We tested the resulting embeddings using both our classification approach, and our similarity detection approach. We calculated the cosine similarity between the representations of the original dataset, *All*, versus three other datasets and plotted the distribution of cosine similarities.

## Results

### Classification task

Our goal is to compare the three representation approaches. The number of features in each approach, which corresponds to the dimensionality of the representations, ranges from one hundred to over one hundred thousand (Table 2).

**Table 2:** Number of features for each representation

| Representation method | Number of features |  |  |
| --- | --- | --- | --- |
|  | Antibody | Cell line | Tissue |
| union representation | 136284 | 180424 | 150681 |
| tf.idf - 1000-kb | 50100 | 40596 | 43912 |
| tf.idf - 500-kb | 28527 | 22783 | 25238 |
| tf.idf - 100-kb | 8262 | 7225 | 7791 |
| region-set2vec embedding | 100 | 100 | 100 |

#### Classification performance

We evaluate the representations by the performance of the classifier trained to identify biological information for each region set. Therefore, for each representation method, we trained an SVM classifier on cell line, antibody, and tissue training sets and report the results on the test sets (Table 3; See Supplementary Material).

**Table 3:** SVM classifier performance

| Representation method | F1 score_SVM |  |  |
| --- | --- | --- | --- |
|  | Antibody | Cell line | Tissue |
| union representation | 0.9424 | **0.9898** | **0.9591** |
| tf.idf - 1000-kb | **0.9468** | 0.9788 | 0.9523 |
| tf.idf - 500-kb | 0.9439 | 0.9613 | 0.9500 |
| tf.idf - 100-kb | 0.9022 | 0.8977 | 0.9568 |
| region-set2vec embedding | 0.9381 | 0.9605 | 0.9091 |

The classifier using the union representation performed well because all of the features in the universe were retained. On a similar note, the representation using tf_idf with 1000-kb also performed well on the antibody classification task, due to the high number of base pairs selected. The region-set2vec embedding performed better than the tf_idf representation with 100-kb on the antibody and cell line classification, but performed the worst on the tissue classification. It seems that the tissue classification worked well even with 100-kb selected in the tf_idf, possibly indicating that the feature selection method found few significant regions signalling the tissue type. The region-set2vec embedding with 100 dimensions performed as well as the other high-dimensional representations on antibody and cell line datasets. In addition, it produces low-dimensional representations that cause the downstream run-time for classification to be much faster (Table 4). On average, training and testing the classifiers on region set embeddings are 2500 times faster than the union representation. We also compared the run-time of building the representation model and the required time to transform a new test dataset to the numerical representations. Despite higher run-time in training the region2vec model, transforming the test dataset occured faster in region-set2vec representation methods. The union and tf_idf representations, a region-set needs to be mapped to the high dimensional space and then use the saved dimension reduction model to reduce the dimension to 100 while region-set embedding is calculated in low dimension (see Supplemental Material).

**Table 4:** SVM training run-time (seconds)

| Representation method | Antibody | Cell line | Tissue |
| --- | --- | --- | --- |
| union representation | 402.60 | 2480.97 | 348.38 |
| tf.idf - 1000-kb | 129.12 | 540.65 | 101.28 |
| tf.idf - 500-kb | 71.90 | 315.59 | 58.76 |
| tf.idf - 100-kb | 21.52 | 92.99 | 17.64 |
| region-set2vec embedding | 0.21 | 0.78 | 0.17 |

Given the vast difference in number of features, we next sought to conduct a fairer performance comparison by restricting the dimensions of all methods to 100. Therefore, we selected the top 100 components using Principal Component Analysis (PCA)^33^. In addition to the linear PCA, we also tried two non-linear kernel PCA appraoches: *rbf* and *polynomial*. We selected the best PCA-kernel using cross-validation score in the same way as for the SVM kernel. The best results on each dataset were achieved by the *linear* kernel for PCA (Table S4, Table S5, and Table S6). Using these 100-dimensional features as inputs to the SVM classifier, the region-set2vec embedding performs the best for all three classification tasks (Table 5). The other methods, now with the same dimensions as the region-set2vec embedding, have reduced performance, showing that the region-set2vec embedding retained the most information in 100 dimensions.

**Table 5:**
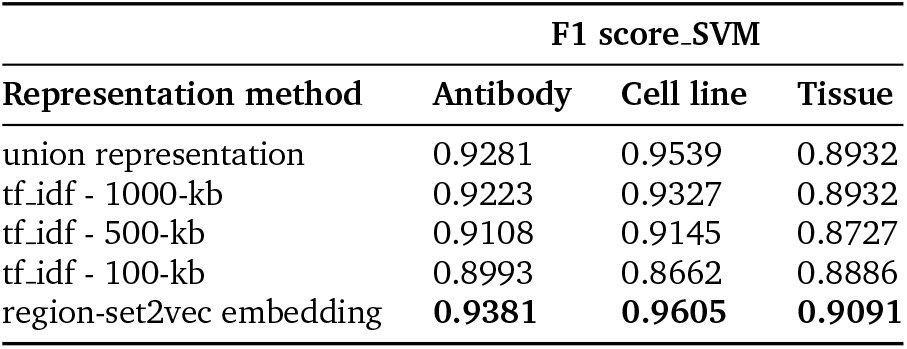
SVM-PCA classifier performance

| F1 score_SVM |  |  |  |
| --- | --- | --- | --- |
| Representation method | Antibody | Cell line | Tissue |
| union representation | 0.9281 | 0.9539 | 0.8932 |
| tf.idf - 1000-kb | 0.9223 | 0.9327 | 0.8932 |
| tf.idf - 500-kb | 0.9108 | 0.9145 | 0.8727 |
| tf.idf - 100-kb | 0.8993 | 0.8662 | 0.8886 |
| region-set2vec embedding | **0.9381** | **0.9605** | **0.9091** |

To evaluate the performance, we used 10-fold crossvalidated paired t-tests. We first split the training data into 10 folds of equal size. In each cross-validation iteration, we compute the difference in performance between classifying the data represented by the union method and the same data represented by the region-set2vec approach. The test shows that the higher performance of region-set2vec is statistically significant for all datasets (p-values< 0.003 for antibody; 0.000036 for cell line; and 0.041 for tissue dataset).

We used the area under the Precision-Recall curve (AUPR), to visualize the performance classification of antibody, cell line, and tissue on different representations (Fig. 3a). Higher values indicate the trained classifier can better distinguish between classes. With region-set2vec embeddings, the trained classifiers achieve the highest area under the curve scores in all three datasets.

**Figure 3:**
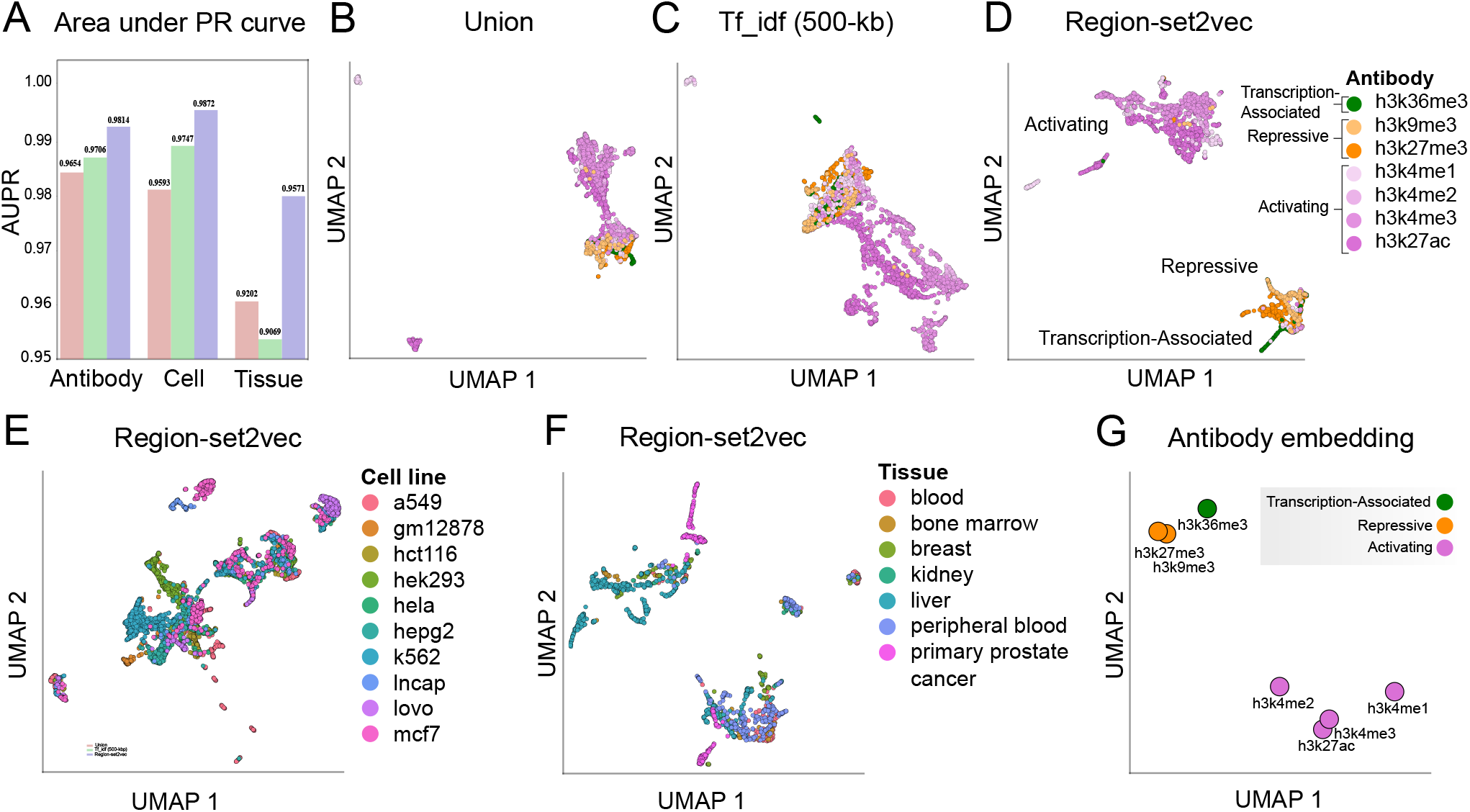
Classification performance and UMAP visualizations of representations. (**a**). Area under the micro-average PR curve for each dataset. (**b**). Antibody dataset represented by union method. (**c**). Tf_idf representation using 500-kb of antibody dataset. Each point represents a region set. (**d**). Region-set2vec embedding of antibody dataset using combined region embeddings trained by word2vec algorithm. (**e**). Region-set2vec representation of cell line dataset. (**f**). Region-set2vec representation of tissue type datasets. Panels a and b show the UMAP visualization of the original data, not the PCA-reduced data. (**g**). A UMAP projection of antibody embeddings annotated by class label. Each point is the combination of all the samples in each class.

#### UMAP visualization

For each of our three datasets, we used UMAP to visualize the representation space (Fig. 3b-g). We selected 100 as the number of neighbors and Euclidean distance as the similarity metric. Since the three tf_idf representations had similar results, we only plot the 500-kb tf_idf representation. Union representation resulted in mapping most of the samples in the antibody dataset into a similar space on the UMAP plot (Fig. 3b). Using tf_idf representation, the samples are more distinctive due to the selection of discriminant features (Fig. 3c). The region-set2vec embeddings show more distinctive, tight clusters (Fig. 3d), and clearly distinguish antibodies associated with repression, activation, and transcription. Similarly, cell lines and tissue types are clustered effectively by the region-set2vec representation (Fig. 3e and 3f).

To represent each type of antibody with a numerical vector and as a single point in 2-d space, we merged all representations of the samples in the dataset using averaging as the combination function. The classes of antibody associated with active gene expression (*h3k4me1, h3k4me2, h3k4me3* and *h3k27ac*) are mapped closer to each other and further away from repressive marks (*h3k27me3* and *h3k9me3*), and transcription-associated (*h3k36me3*) antibody classes (Fig. 3g). This plot indicates that our learned representations retain biological information like antibody type without using this information in the training phase.

### Similarity detection task

As a second independent evaluation of our region embeddings, we employed a similarity detection task. In this task, we simulated perturbations to a given BED file at a range of pre-specified rates, and then examined how the differences in embedding reflect the known perturbation rates.

#### UMAP plots

To visualize how the embeddings reflect the perturbation, we used UMAP plot with 100 neighbours, Cosine as the distance metric. The union and tf_idf representations are randomly distributed in the UMAP 2-dimensional space, despite the perturbation rate ranging in increments of 10 percent (Fig. 4a,b). In contrast, the UMAP visualization of the region-set2vec embeddings showed a gradual deviation as the percentage of the perturbation increased (Fig. 4c). This result indicates that, at least for the purpose of UMAP visualization, the region-set2vec embeddings are able to reflect quantitative similarity among region sets, whereas the other methods are not. We also tested other UMAP parameters with a sensitivity analysis and found that using 100 neighbors, 0.01 as the minimum distance and Cosine as the distance metric (Fig. S1a-d) produced sensible and robust visualizations (see Supplemental Material). We also found that these results are consistent when perturbing files with *add*, *drop*, and *shift* independently (Fig. S2).

**Figure 4:**
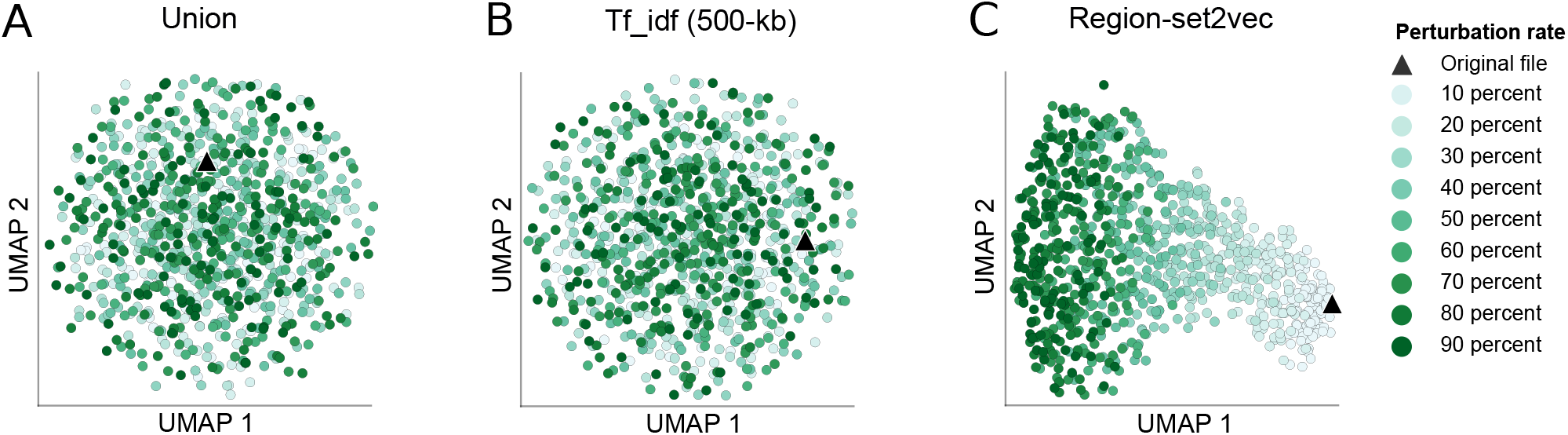
UMAP visualization of the simulated dataset with different rates of perturbation using 3 representation methods. (**a**). Union representation (**b**). Tf_idf representation with 500k important base pairs (**c**). Region-set2vec representation.

### Peak thresholds subsampling task

To investigate the robustness of different representations, we ran the classification task on the original cell line dataset and three truncated datasets generated by selecting the top 75%, 50%, and 25% of the regions in each bed file based on the signal values. As expected, the performance of the classification task decreased as the files were more truncated; however, the region-set embedding representation method was least sensitive to the truncation (Fig. 5a). The region-set embedding representation performance is still >94% even when considering only the top 25% of peaks, compared to <84% for the Tf_idf approach. This indicates that this approach preserves the information of the missing regions by learning the embeddigns based on the context.

**Figure 5:**
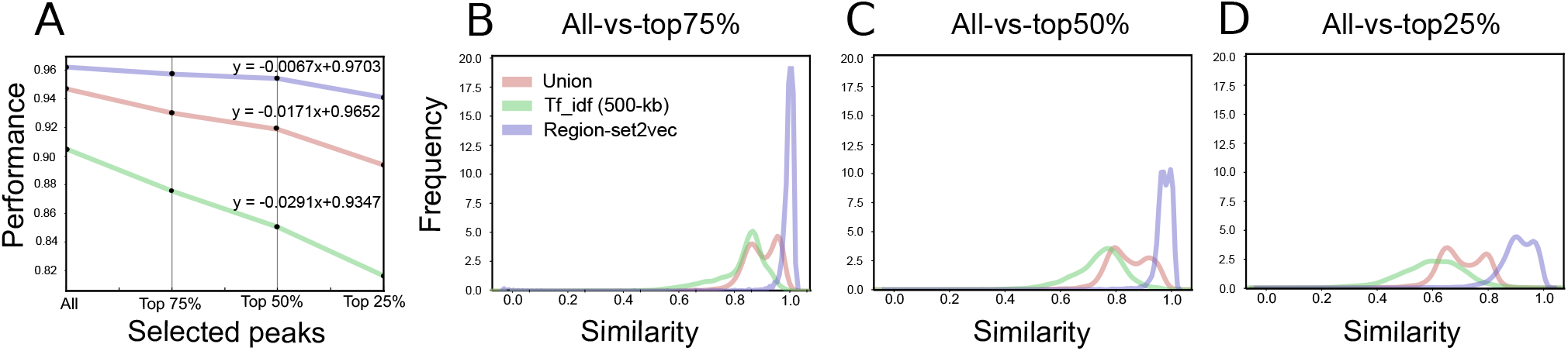
Region set embeddings are more robust to peak calling threshold. (**a**). Sensitivity of different representation to selected peaks based on signal values. Distribution of similarity between the original region sets and (**b**). top 75% of the selected peaks, (**c**). top 50% of the selected peaks, and (**d**). top 25% of the selected peaks.

We also evaluated the robustness of the representation by calculating the Cosine similarity between the representations of the original dataset and each truncated dataset. The distributions of Cosine similarities show the region-set2vec vector based on abbreviated data are most similar to the vectors based on the full data for all levels of truncation (Fig. 5b-d). This result again confirms that the region-set2vec approach retains the most information in the face of subsetted data.

### Application of region-set2vec to single-cell data

Satisfied that region-set2vec could capture important relationships, we next sought to apply this approach as a proof-of-concept to discriminate cell types within single-cell ATAC-seq (scATAC) data. We used a simulated scATAC bone marrow dataset composed of six known tissues derived from an original set of bulk FACS-sorted data^34^. We processed this feature matrix and shuffled 25 times and evaluated with the following hyperparameters: 100 dimensions, a minimum count of 2, 12 neighbors, and a window size of 250. Our adapted word2vec approach successfully separated each cell type into distinct clusters, similar to the previous best-performing candidates^34^(Fig. 6).

**Figure 6:**
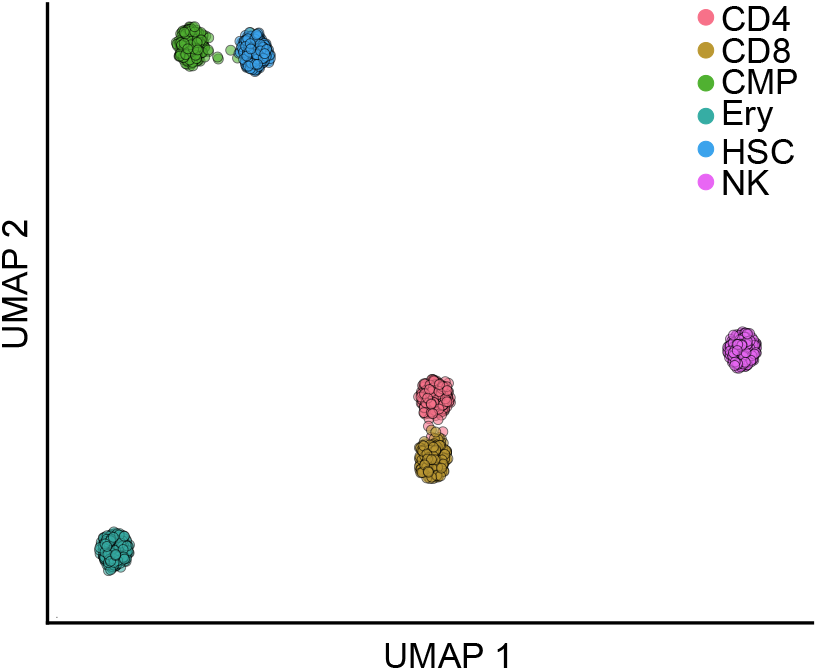
UMAP visualization of simulated scATAC bone marrow data with 2500 fragments per cell^34^. Cells are colored by cell type.

## Discussion and Conclusion

In this paper, we proposed three feature selection methods to represent genomic regions sets: union representation, tf_idf representation, region-set2vec embedding, which we evaluated via classification, similarity detection, and peak threshold robustness tasks. In the classification task, the region-set2vec method underperformed against other higher-dimensional vectors; however, after reducing the dimensionality of all methods to the same dimensions, the region-set2vec method outperformed all others significantly. Region-set2vec vectors also resulted in better visual class separation, and further showed distinction between, activating, repressive and transcription marks. This distinction was previously noted with the Avocado approach^23^, but Avocado considered the biological annotation in the training phase of the antibody embeddings; in contrast, our embeddings are learned solely from co-occurrences of regions within unannotated region sets.

In the similarity detection task, we showed that known interval-based similarity is better projected using region-set2vec representations. One possible explanation for this result is that the embedding vector for each region is trained in a way that preserves the information of the context (*i.e*., BED file). Because the vector embedding considers the co-occurrence of regions in the same file as a context, when some of the regions are dropped or shifted in a perturbed file, the information about these regions is conserved in the embedding of the remaining regions and consequently in the final representation. In contrast, the other methods consider regions as independent features, and the representation does not reflect the relationship among features. As a result, dropping, adding, or shifting the regions could cause abrupt transformation between the original file and the perturbed files. Embeddings that retain information across regions, therefore, appear to be less susceptible to such random perturbations. Along similar lines, our peak threshold robustness test showed that the region-set2vec vectors retained the most similarity to the original vectors, achieving high similar scores even when only 25% of the data is considered.

Overall, we demonstrated the feasibility of using NLP techniques to represent genomic region sets data in new ways that will drive analysis methods in the future. In the future, it may be possible to improve the quality of the region-set2vec embedding by optimizing the hyperparameters. For example, here we arbitrarily chose 100-dimensional vectors, but it may be that a higher (or lower) number is optimal. In addition, we have several ideas for addressing the problem for building the universe of regions. We are now working to address this upstream problem and hope to have a general solution in the future. Furthermore, we have used a relatively limited collection of BED files, which can be extended with additional data sources. Applying the method on other source of data and mutated regions are the potential future directions. Altogether, our results indicate that low-dimensional representations of region sets built using nothing more than unsupervised collections of region set data can be an effective approach to build biologically meaningful vector representations.

## Acknowledgements

We would like to thank Tessa Danehy for her valuable comments.

## Supplemental Materials

### Term frequency–inverse document frequency

Term frequency-inverse document frequency (TFIDF)^24^ measures how important a term is to a document in a corpus. It is often used to provide weighted representations of documents for information retrieval. Such metric can be factored into term-frequency (TF) and inverse document frequency (IDF). Binary TF measures the existence of a term in a document, and is given in the following equation:

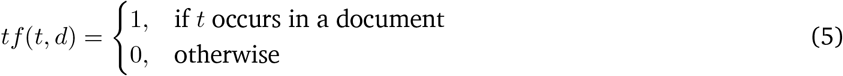

where *t* is the term and *d* is the document. IDF is defined as the logarithm of the inverse fraction of the number of documents that contain the term *t*, as expressed in the following equation:

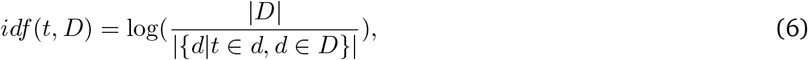

where *D* represents the corpus, i.e. a set of documents *d*; the denominator is the number of documents in which term *t* appears; and the |·| denotes set size. IDF assigns larger values to terms that occur less frequently across documents in the corpus. It is usually employed to find terms that do not occur frequently but are considered meaningful in documents.

Finally, combining TF and IDF scores, the tf_idf value is calculated in Equation 7 as:

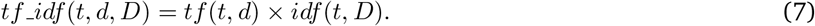

Considering base pairs as terms, we will use this method to score each base pair in a region sets and select top-scoring regions for further process.

### Word Embedding

Traditional approaches in NLP represented text as a sparse vector with the length corresponding to the size of the vocabulary. Named one-hot representation, this method creates a vector of zeros and ones representing word occurrence in the vocabulary. Alternatively, the values of the vector can be the number of word occurrences in the document. Distributed word representation, generally known as word embedding, is used to solve the problems of high dimensionality and sparsity in the representation. In this representation, each word is described by the surrounding context^35^ which contains semantic and syntactic information about the word. A language modeling task is employed to construct such representations. Distributed representation learning is first introduced by Hinton^36^ and is developed as a language modeling concept by Bengio^37^.

Word2vec^11^ is a series of methods to represent a word using a numerical vector. The main idea of building this representation is to express a word by its context by training a shallow neural network. Each row of the weight matrix of the neural network represents a word vector. The neural network takes in a one-hot encoded word as its input, then passes it through hidden layers to get an output that predicts the word, given its context. The weights of the hidden layer are updated if the words share the same context. After the training, the hidden layer weights represent a vector for each word. In our approach, we adapt word2vec to genomic region sets and use the trained model to extract region embedding vectors, which we then combine to build region set embeddings.

### SVM and PCA kernel analysis

We selected the kernel for the SVM classifier based on the cross-validation results on the training dataset. We split the training data into 10 folds to choose the best hyperparameters using the cross validation score. *Linear, rbf*, and *polynomial* kernel were employed and linear kernel was chosen due to the higher average of the scores across all folds. The same kernels were used for PCA dimension reduction and *Linear* kernel achieved the best results (Table S4, Table S5, and Table S6).

### Run-time analysis

There are two steps in learning representations for a dataset. 1) building the representation models on the training data and 2) transforming the new test set to the target representations. Regarding run-time analysis, in the model building phase on the training data, word2vec does require more time to train the neural network, but after the training, we can simply use and update the model. The training time depends on hyperparameters such as dimension size, number of documents, and vocabulary size. For example, on average it takes 45 minutes for 50-dimension vectors and 140 minutes for 100-dimension vectors on our data. The union and tf_idf representation use 100 files to build the feature set and in fact there is no training. Therefore, the run-time to build these models are faster than training the word2vec model (Table S7).

However, in the second step for transforming new test dataset, for the union and tf_idf representations, a region-set needs to be mapped to the high dimensional space and then use the saved PCA model to reduce the dimension to 100 while region-set embedding is calculated in low dimension. Transforming a dataset of 695 files to the representation vectors takes around 1 minute for the region-set2vec model and around 2.5 minutes to transform and reduce the dimension for union representation (Table S8).

### UMAP parameter analysis

To study the effect of UMAP dimension reduction parameters, we used different values to generate various two-dimension plots (Table S9). We found that using 100 (50, 100, 150, and 200 were tested) neighbors is a good choice to produce sensible and robust visualizations when changing the other hyperparameter. The distinction between the final plots for different values of minimum distance was negligible. With the 100 neighbors in the UMAP configuration, we chose Euclidean, Cosine, Jaccard, and Dice as the similarity metrics with Jaccard and Dice used specifically for the binary representations. We observed that 2-cluster structure is evident in the Euclidean distance plots of both the union and tf_idf representations, although in the Jaccard and Dice plots, this structure still exists but it is less obvious. We conjectured that the 2-cluster structure shown in the Euclidean distance plots is due to the following two reasons. Since a binary representation is robust to small perturbations, such as a small percentage of dropping or shifting, after the perturbation rate exceeds a certain level, the Euclidean distance will change significantly, and the UMAP captures the small change and big change as two distinct clusters in the reduced dimension space. Moreover, since the other distance metrics are normalized versions of the raw change with a normalization constant not reflecting the perturbation rates, their plots cannot produce distinctive clusters. Compared to the binary representations, the region-set2vec representation is able to reflect gradual changes under both the Euclidean and Cosine distance metrics (Fig. S1a-d).

To investigate the effect of each aspect of perturbation on the final representation, we generated three different datasets by varying one aspect of perturbation at a time, add, drop, and shift. Then we plot the new datasets for exploration. First, we randomly add regions to the sample file to create perturbation (Fig. S2a). Dropping regions from the file had the same effect on the final representations (Fig. S2b). By shifting the regions some of the regions drop out of the regions in the universe and some overlap the regions in the universe of possible regions (Fig. S2c). Increasing the rate of perturbation on this aspect gradually change the region-set embeddings while there are no obvious pattern in other representations. This indicates that the similarity between the original file and the perturbed file is preserved in the region-set2vec representation rather than the other methods on every aspect of perturbation.

### Supplementary figures

**Figure S1:**
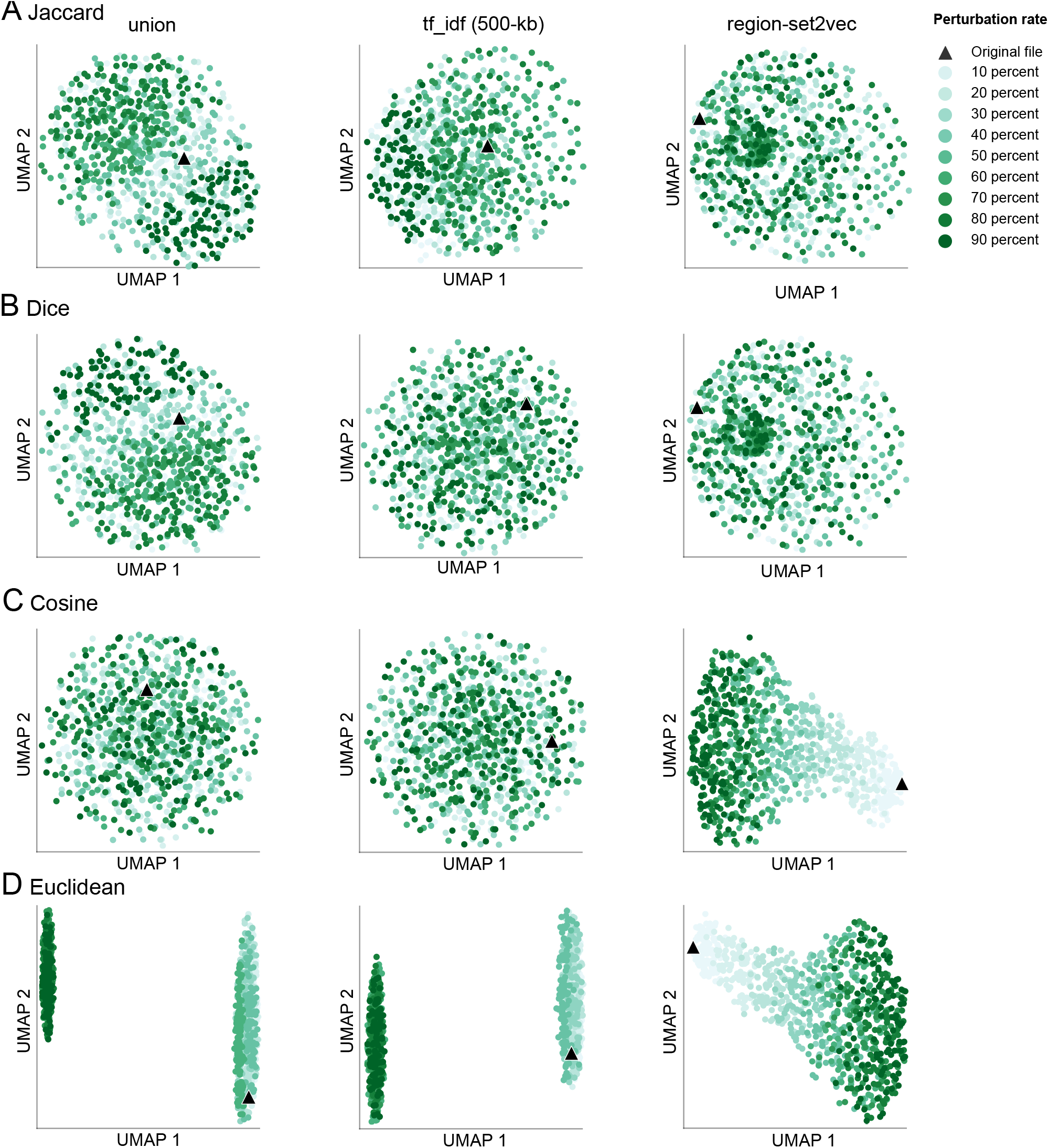
UMAP visualization with different similarity metrics for simulated dataset using three representation methods. Jaccard (**b**). Dice (**c**). Cosine (reproduced from Figure 4) (**d**). Euclidean. Jaccard and Dice distance were poor metrics for all three representations, while cosine and Euclidean distance show that region-set2vec has a better ability to capture similarity.

**Figure S2:**
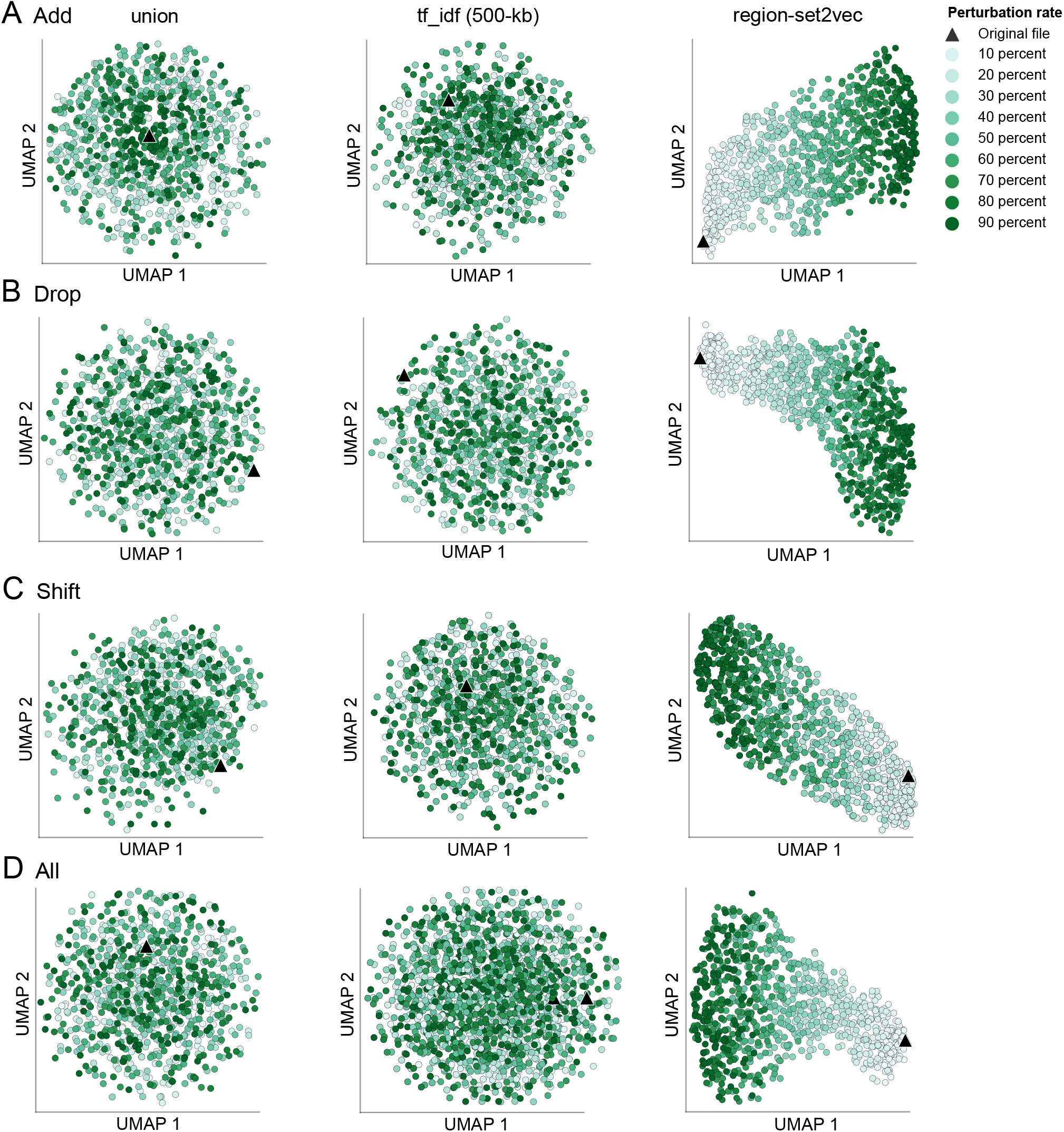
UMAP visualization of the simulated dataset with perturbations made individually on each of the three representation methods. Cosine is used as the distance metric in UMAP method. (**a**). Adding regions to each file. (**b**). Dropping regions from each file. (**c**). Shifting regions in each file. (**d**). Combination of all three types of perturbation (reproduced from Figure 4). When adding and shifting regions, it appears that union and tf_idf representations do not capture similarity, as all of the points spread out without any pattern. However, the region-set2vec representation indicates more perturbed files are more distant than less perturbed files. When only dropping regions, the UMAP plots look similar to when all perturbations are present, because dropping regions is the only perturbation guaranteed to make changes to the vector representations.

### Supplementary tables

**Table S1:** Union representation properties

|  | Antibody | Cell line | Tissue |
| --- | --- | --- | --- |
| # regions | 136,284 | 180,425 | 150,681 |
| median region length | 393 | 462 | 428 |
| mean region length | 761 | 1,057 | 801 |

**Table S2:** SVM-PCA classifier performance robustness test. We tested 20 different universes created from different sets of 100 random BED files to confirm that the random selection did not affect the classifier performance.

|  | Antibody | Cell line | Tissue |
| --- | --- | --- | --- |
| 1 | 0.9209 | 0.9547 | 0.8932 |
| 2 | 0.9180 | 0.9518 | 0.9045 |
| 3 | 0.9194 | 0.9598 | 0.9023 |
| 4 | 0.9122 | 0.9591 | 0.9000 |
| 5 | 0.9165 | 0.9591 | 0.9205 |
| 6 | 0.9122 | 0.9572 | 0.9023 |
| 7 | 0.9180 | 0.9561 | 0.8932 |
| 8 | 0.9122 | 0.9581 | 0.9000 |
| 9 | 0.9165 | 0.9494 | 0.9114 |
| 10 | 0.9180 | 0.9568 | 0.8932 |
| 11 | 0.9151 | 0.9477 | 0.9114 |
| 12 | 0.9137 | 0.9485 | 0.9068 |
| 13 | 0.9094 | 0.9522 | 0.8977 |
| 14 | 0.9209 | 0.9583 | 0.9136 |
| 15 | 0.9137 | 0.9575 | 0.9091 |
| 16 | 0.9180 | 0.9551 | 0.9136 |
| 17 | 0.9122 | 0.9498 | 0.9045 |
| 18 | 0.9089 | 0.9527 | 0.9091 |
| 19 | 0.9201 | 0.9469 | 0.9000 |
| 20 | 0.9187 | 0.9569 | 0.9023 |

**Table S3:** Comparison on SVM classifier performance on averaging and doc2vec combination function

| Representation method | Antibody |
| --- | --- |
| union representation | 0.9317 |
| tf_idf - 1000-kb | 0.9101 |
| tf_idf - 500-kb | 0.9317 |
| tf_idf - 100-kb | 0.8885 |
| region-set2vec embedding-averaging | **0.9568** |
| doc2vec | 0.5606 |

**Table S4:**
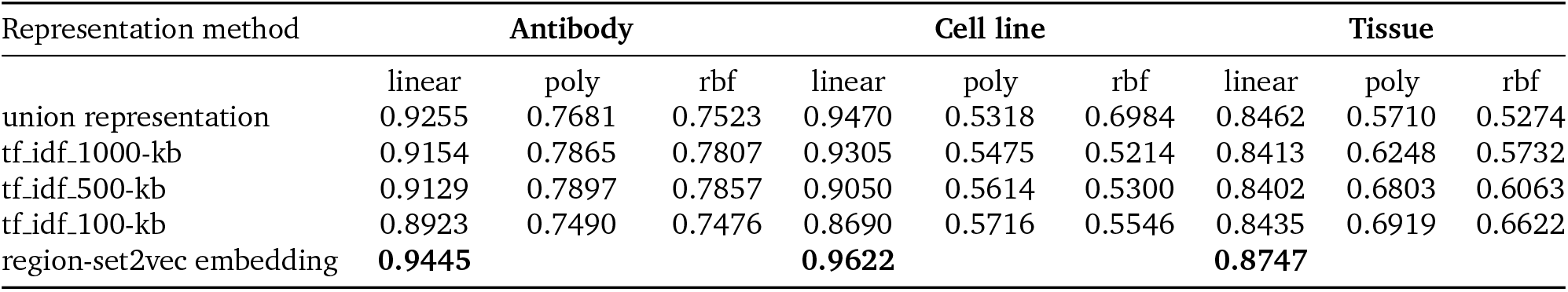
SVM classifier performance with linear kernel

**Table S5:** SVM classifier performance with RBF kernel

| Representation method | Antibody |  |  | Cell line |  |  | Tissue |  |  |
| --- | --- | --- | --- | --- | --- | --- | --- | --- | --- |
|  | linear | poly | rbf | linear | poly | rbf | linear | poly | rbf |
| union representation | 0.8592 | 0.8451 | 0.8606 | 0.5111 | 0.4869 | 0.6850 | 0.7729 | 0.7696 | 0.7746 |
| tf_idf_1000-kb | 0.8174 | 0.8088 | 0.8196 | 0.5370 | 0.4674 | 0.5527 | 0.7740 | 0.7347 | 0.7757 |
| tf_idf_500-kb | 0.8088 | 0.8027 | 0.8109 | 0.5171 | 0.4194 | 0.5381 | 0.7751 | 0.7185 | 0.7768 |
| tf_idf_100-kb | 0.7976 | 0.7612 | 0.8016 | 0.4892 | 0.3964 | 0.5567 | 0.7817 | 0.7615 | 0.7857 |
| region-set2vec embedding | **0.9237** |  |  | **0.9236** |  |  | **0.8328** |  |  |

**Table S6:** SVM classifier performance with polynomial kernel

| Representation method | Antibody |  |  | Cell line |  |  | Tissue |  |  |
| --- | --- | --- | --- | --- | --- | --- | --- | --- | --- |
|  | linear | poly | rbf | linear | poly | rbf | linear | poly | rbf |
| union representation | 0.8167 | 0.8027 | 0.8239 | 0.4829 | 0.5495 | 0.7049 | 0.6253 | 0.6174 | 0.6275 |
| tf_idf_1000-kb | 0.8023 | 0.7911 | 0.8077 | 0.7362 | 0.6946 | 0.7425 | 0.5973 | 0.5911 | 0.6484 |
| tf_idf_500-kb | 0.8023 | 0.7875 | 0.8073 | 0.7232 | 0.6798 | 0.7342 | 0.6007 | 0.5827 | 0.6669 |
| tf_idf_100-kb | 0.8063 | 0.7623 | 0.8142 | 0.7176 | 0.6593 | 0.7246 | 0.6633 | 0.5673 | 0.7071 |
| region-set2vec embedding | **0.8790** |  |  | **0.9025** |  |  | **0.7494** |  |  |

**Table S7:** Run-time to build the representation models

| Representation method | Building models |
| --- | --- |
| union representation | 00:03:00 |
| tf_idf | 00:09:00 |
| region-set2vec embedding | 02:20:00 |

**Table S8:** Run-time to transform test dataset to representations for downstream tasks

| Representation method | Vectorization | PCA | Total |
| --- | --- | --- | --- |
| union representation | 0:01:48 | 0:00:27 | 0:02:15 |
| tf_idf_1000-kb | 0:00:58 | 0:00:12 | 0:01:10 |
| tf_idf_500-kb | 0:00:50 | 0:00:08 | 0:00:58 |
| tf_idf_100-kb | 0:00:15 | 0:00:03 | 0:00:18 |
| region-set2vec embedding | 0:01:04 |  | 0:01:04 |

**Table S9:** Values of the hyperparameters

| Hyperparameter | Values |
| --- | --- |
| number of neighbors | 50, 100, 150, 200 |
| minimum distance | 0.1, 0.01, 0.05 |
| distance metric | Euclidean, Cosine, Jaccard, Dice |

